# Shared transcriptional responses to con- and heterospecific behavioral antagonists in a wild songbird

**DOI:** 10.1101/795757

**Authors:** Matthew I. M. Louder, Michael Lafayette, Amber A. Louder, Floria M. K. Uy, Christopher N. Balakrishnan, Ken Yasukawa, Mark E. Hauber

**Affiliations:** Department of Evolution, Ecology, and Behavior, School of Integrative Biology, University of Illinois at Urbana-Champaign, USA; International Research Center for Neurointelligence, University of Tokyo; Department of Biology, Beloit College, Beloit, WI, USA; Department of Neurobiology and Behavior, Cornell University, Ithaca, NY, USA; Department of Biology, East Carolina University, Greenville, NC, USA

**Keywords:** auditory recognition, biomarker, blood, host-parasite interactions, RNA sequencing, territory defense

## Abstract

The recognition of and differential responses to salient stimuli are among the main drivers of behavioral plasticity, yet, how animals evolve and modulate functional responses to novel classes of antagonistic stimuli remain poorly understood. We studied free-living male red-winged blackbirds (*Agelaius phoeniceus*) to test whether gene expression responses in blood are distinct or shared between patterns of aggressive behavioral responses directed at simulated conspecific versus heterospecific intruders. In this species, males defend territories against conspecific males and respond aggressively to female brown-headed cowbirds (*Molothrus ater*), a brood parasite that commonly lays eggs in blackbird nests. Both conspecific songs and parasitic calls elicited aggressive responses from focal subjects and caused a downregulation in genes associated with immune system response, relative to control calls of a second, harmless heterospecific species. In turn, only the conspecific song treatment elicited an increase in singing behavior and an upregulation of genes associated with metabolic processes relative to the two heterospecific calls. Our results suggest that aspects of antagonistic responses to both conspecifics and brood parasites can be based on similar physiological responses, suggestive of shared molecular and behavioral pathways involved in the recognition and reaction to both evolutionarily old and new enemies.

## Introduction

Faced with a suite of relevant and irrelevant stimuli, animals must rapidly perceive, process, and decide whether and how behaviorally to respond to diverse cues ^1^. Upon the recognition of salient stimuli, for example, a cascade of neurophysiological and motor responses can be engaged to facilitate and modulate a reaction ^2^. Yet, how recognition systems adapt and enable responses to evolutionary novel stimuli remain poorly understood ^3^. In particular, the timely and accurate recognition of heterospecific antagonists, such as parasites, predators, or competitors, typically have substantial fitness advantages ^4^. Can evolutionarily established behavioral and physiological responses to conspecific competitors be co-opted in parallel to adaptively respond to heterospecific antagonists?

Hosts of avian brood parasites pay the costs of raising unrelated young, often with the additional expense of losing some or all of their own offspring ^5^. Many hosts have evolved to combat brood parasitism by attacking adult parasites, abandoning parasitized nests, and/or rejecting parasitic offspring ^6^. Anti-parasitic defenses can take categorically different responses from competitive (e.g. territorial defense with song and overt aggression against same-sex conspecific intruders) or anti-predatory behaviors, and can even be evoked by a partial suite of (visual or auditory only) sensory cues ^7,8^.

What constitutes the physiological basis of anti-parasite responses remains largely unknown in avian host-parasite systems ^9^. For example, recent work in wild-caught juvenile male red-winged blackbirds (*Agelaius phoeniceus*; hereafter: redwings) found no differences in immediate early gene (IEG) expression levels within the auditory forebrain in response to the calls of adult female brood parasites (brown-headed cowbirds *Molothrus ater*; hereafter: cowbirds) versus a harmless control species (mourning dove *Zenaida macroura*; hereafter: doves), whereas responses were stronger to conspecific adult female calls ^10^. In the wild, however, juvenile redwings do not hold territories and may not (yet) have been exposed to parasitism by cowbirds. In contrast, breeding adult male redwings are well known for territory defense and anti-brood parasite aggression ^11^ (Fig. 1). The species-typical “conk-a-ree” song is a long-range “keep-out” signal ^12,13^; and is the most common conspecific territory defense display of this species ^14^. Male redwings also respond strongly to parasitic female cowbirds by approaching and attacking the parasites ^15^, but they respond little to harmless sympatric species. Therefore, we set out to test whether antagonistic responses to conspecifics and parasites involve distinct or shared behavioral and physiological responses in freely-behaving adult male redwings on their breeding territories.

**Fig. 1.**
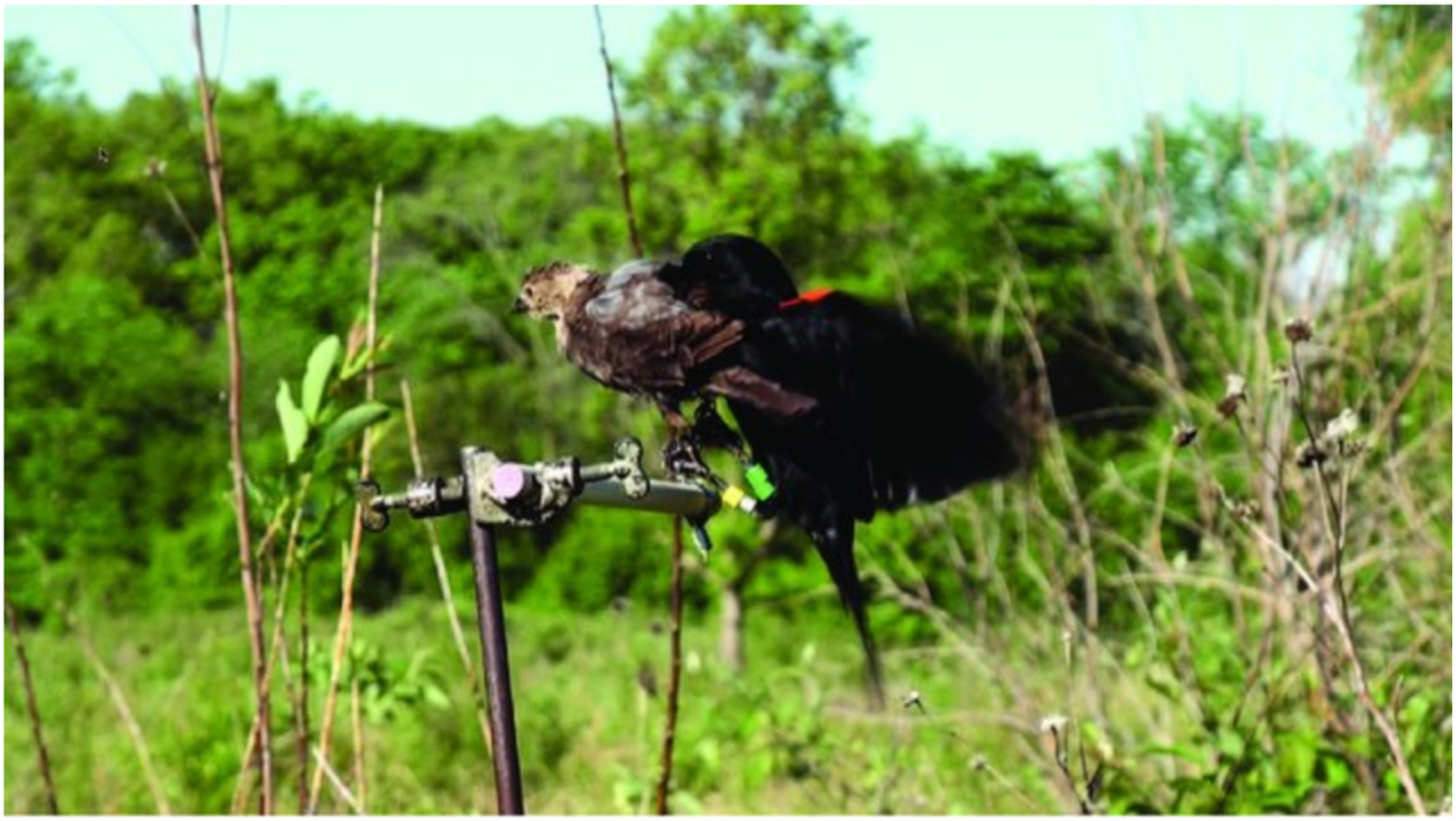
Male red-winged blackbird responding to model presentation of a stuffed female brown-headed cowbird. Photo credit: K. Yasukawa.

Linking evolutionarily relevant and ecologically salient stimuli with their proximate recognition system responses has been difficult in wild animals. This is because, beyond the well-known complexities of field work, lethal collection is required to sample neural tissues during or following the undertaking of a recognition task. Furthermore, such terminal collection prohibits repeated contrasts or ontogenetic comparisons within the same subjects. Alternatively, non-lethal neuroimaging-based techniques, including functional Magnetic Resonance Imaging or micro Positron Emission Tomography, require that subjects become captive during or soon after the recognition task ^16,17^.

Transcriptomic analyses have become a critical tool to analyze functionally relevant cell types and tissues both in model species and, increasingly in non-model species collected from the wild ^18,19^. Moreover, non-terminal collection, such as gene expression within peripheral, whole blood of birds, may reveal functional parallels in the recognition of salient stimuli. For example, a con- vs. heterospecific acoustic playback paradigm in captive zebra finches (*Taeniopygia guttata*) found that gene expression patterns correlated for a subset of genes between the auditory forebrain and in peripheral blood ^20^.

Here we presented playback stimuli of unfamiliar conspecific, parasitic heterospecific, and control heterospecific vocalizations to free-living adult male redwings on their breeding territories, recorded their behavioral responses, then caught them to collect blood samples, and assessed peripheral gene expression patterns. Given the well-known behavioral repertoire of territorial male redwings ^11^, we expected them to respond to playback of conspecific song by increasing their own singing and approaching and remaining in proximity to the playback speaker when compared to playback of a harmless heterospecific song. We expected them to approach and maintain proximity to playback of parasite calls, but not to increase singing, in comparison to the harmless heterospecific. We also predicted distinct and parallel gene activation and behavioral patterns in response to conspecific vs. parasitic stimuli relative to controls.

## Results

### Behavioral response to playbacks

We performed playbacks to 34 territorial male red-wings. Whether or not a subject was caught within 30 min. had no impact on its response behaviors statistically (all *z* > −1.9, *p* ≥ 0.05), nor were there differences in behavioral responses between the 1^st^ and 3^rd^ playback trial segment (see Methods), therefore behavioral data from all subjects and territories in response to the playback paradigm were analyzed statistically (*n*_redwing_ = 11, *n*_cowbird_ = 12, *n*_dove_ = 12). However, transcriptomic data were only available for subjects caught within 30 min of its respective playback set’s delivery (see below for sample sizes). Generalized linear mixed models, with playback type as the independent predictor, revealed that male redwings responses were significantly variable during active playback periods: number of songs *z* = −2.82, *p* = 0.005, number of approaches *z* = −2.50, *p* = 0.01, and nearest distance to the playback speaker (*z* = −2.10, *p* = 0.04), but not in the time spent near the playback speaker (*z* = −0.90, *p* = 0.17) (Fig. 2). Tukey-corrected post-hoc analyses revealed that subjects sang significantly more in response to Redwing than Cowbird (*p* = 0.02) or Dove (*p* = 0.02) playbacks and did not differ significantly in songs in response to Cowbird and Dove playbacks (p = 0.99). Territorial male redwings approached more frequently to Redwing versus Dove (*p* = 0.005) and Cowbird versus Dove playbacks (*p* = 0.03), but did not differ between Redwing versus Cowbird playback (p = 0.79). Territorial male redwings approached more closely to Cowbird than Dove playbacks (*p* = 0.003), but did not differ in response to Redwing vs. Dove (*p* = 0.10), nor to Redwing vs. Cowbird playbacks (*p* = 0.38).

**Fig. 2.**
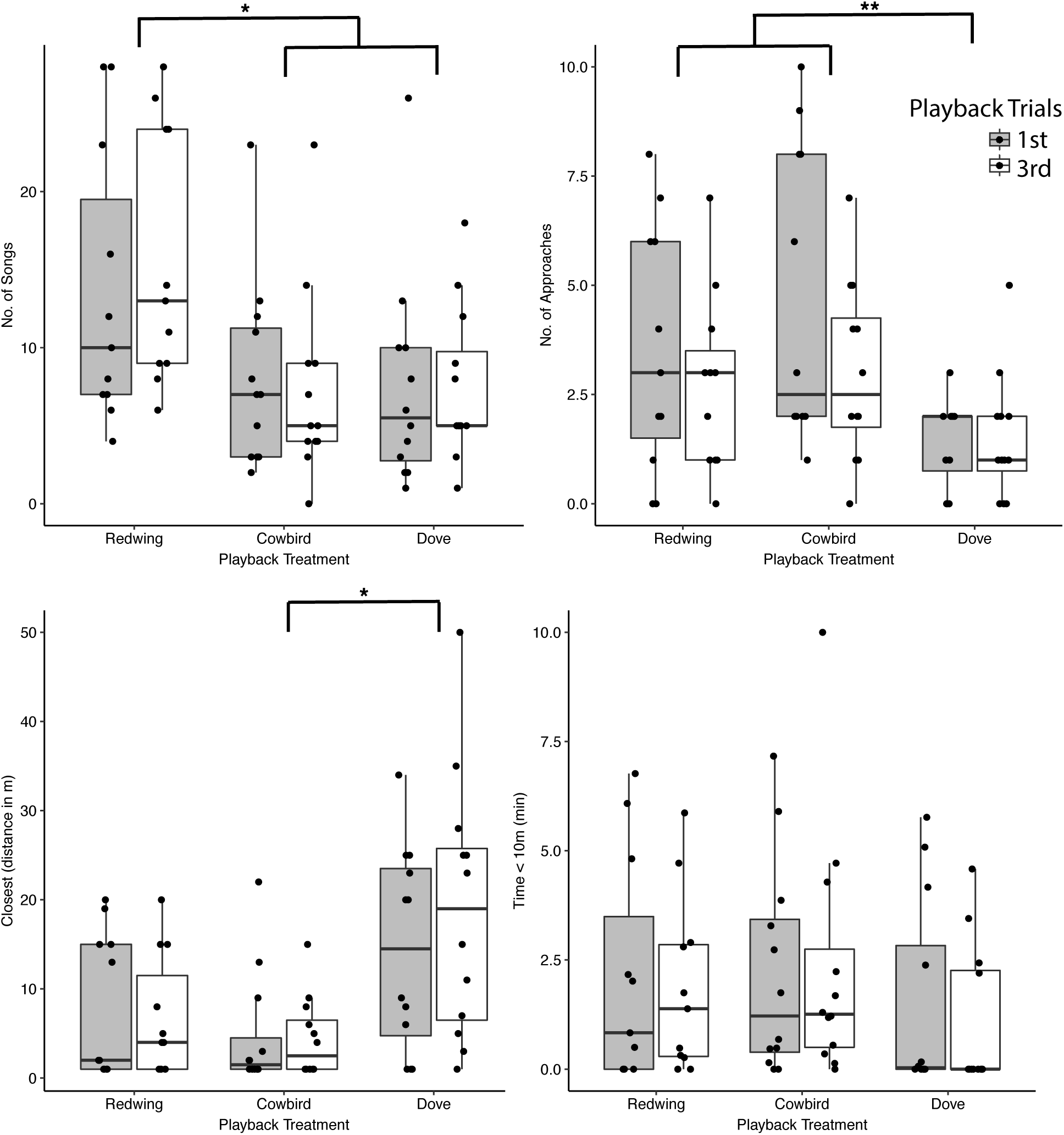
Boxplots depicting the behavioral responses to playback type for male red-winged blackbirds in the 1^st^ (grey boxes) and 3^rd^ (white boxes) trials. Stars denote *:*p* < 0.05 and **: *p* < 0.01 between indicated groups.

### Gene expression

We were able to extract enough RNA from the blood of 21 males upon experimental presentations of playback treatments (*n*_redwing_ = 6, *n*_cowbird_ = 8, *n*_dove_ = 7). For these samples, we sequenced an average of 15.9 million reads per sample (range = 11.7 – 19.1 million reads).

We first tested for differential gene expression for 20 candidate biomarker genes, previously identified from peripheral (whole blood) in Louder et al. ^20^. This analysis yielded no significant differentially expressed genes in response to conspecific songs versus dove coo playback treatments (all Bonferroni corrected *p*-values > 0.50).

After filtering for lowly expressed genes, we then incorporated the read counts of 7202 genes for co-expression analysis (WGCNA). Two modules were significantly correlated with playback treatments (Figs. 3 and 4), the “dark-turquoise” module included upregulated genes in response to Redwing song treatment (*r* = 0.52, *p* = 0.01) and the “yellow” module included genes downregulation in response to both Redwing songs and Cowbird chatter relative to Dove coo (*r* = 0.64, *p* = 0.002).

**Fig. 3.**
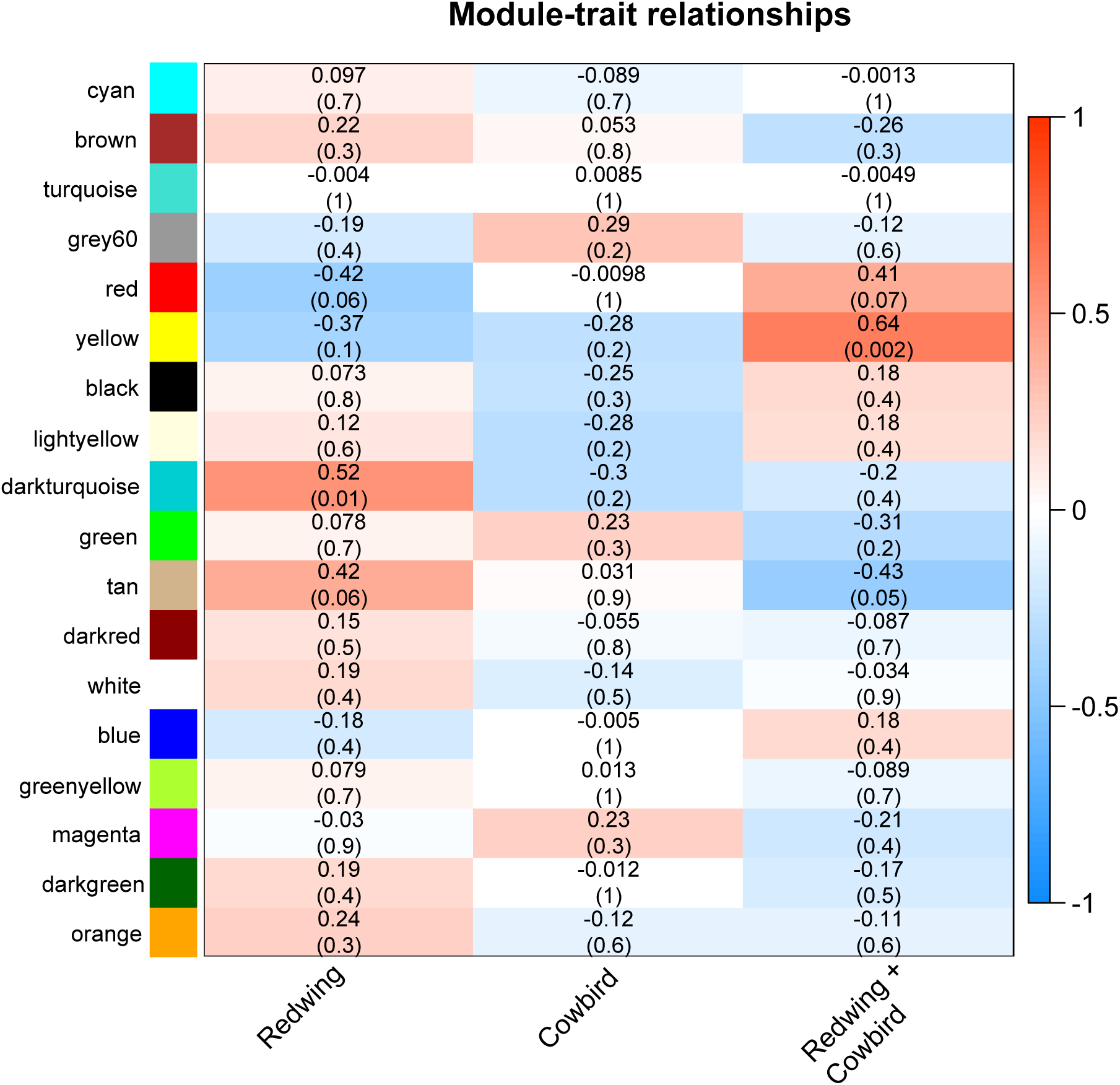
Statistical associations between expression profiles of each WGCNA reconstructed modules and the playback groups. Presented are correlation coefficients and associated *p*-values (within brackets).

**Fig. 4.**
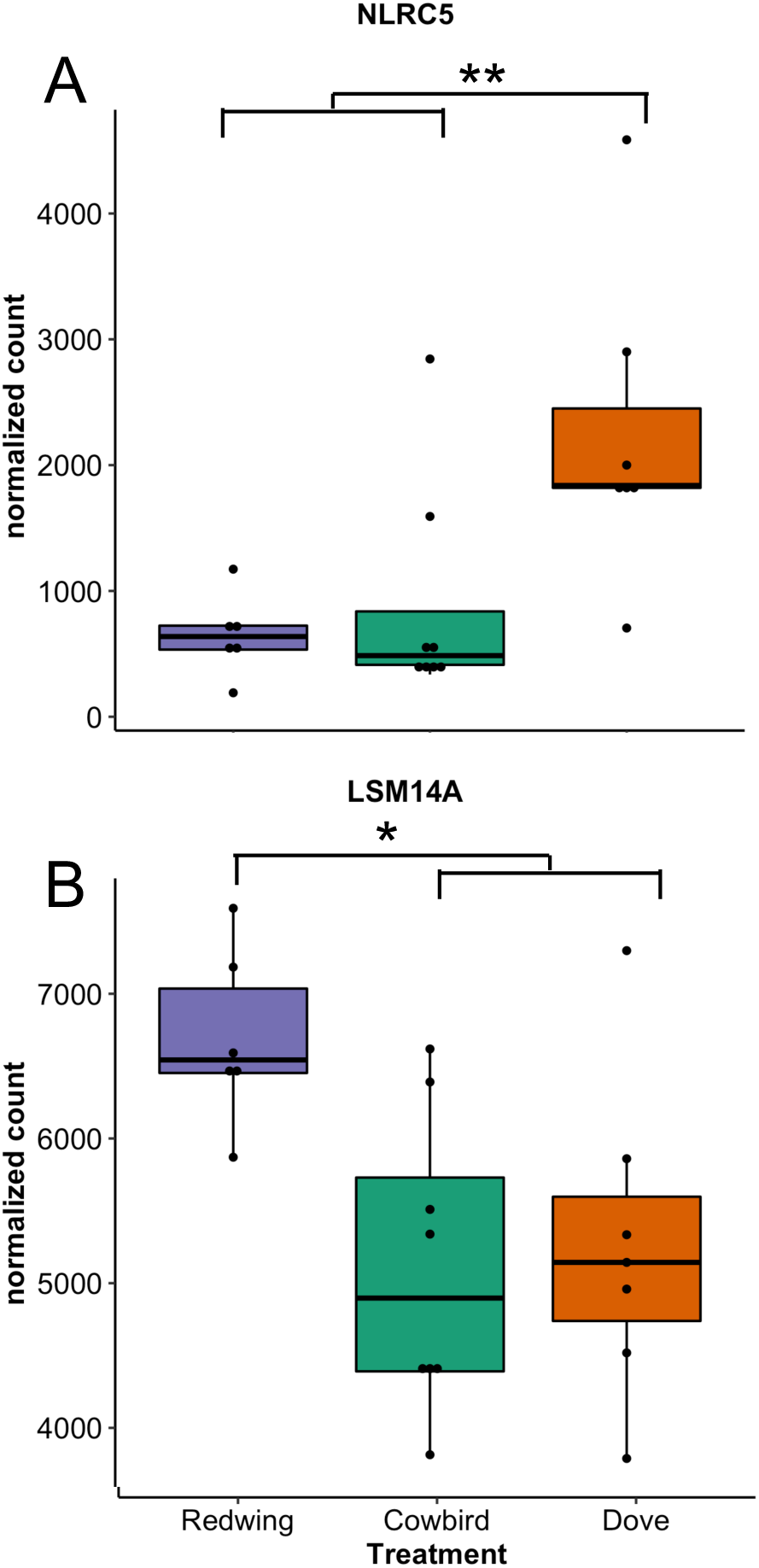
The top gene’s expression profiles from each of the (A) the “yellow” module, which includes genes significantly downregulated in song and chatter treatments relative to dove coo and (B) the “dark-turquoise” module, which includes genes associated with an upregulation in response to the red-winged blackbird song treatment. Stars denote *: *p* < 0.05 and **: *p* < 0.01 between indicated groups.

Using a rank-ordered approach, we identified an enrichment for gene ontology (GO) terms for each significant module. For the “dark-turquoise” module, correlated with a response to Redwing song playback, we found significant GO terms associated with the metabolism, regulation of gene expression, and protein ubiquitination (Table 1). For the “yellow” module, correlated with a response to both Redwing song and Cowbird playback, we find significant GO terms associated with immune system response, such as defense responses to virus and cytokine-mediated pathways (Table 1).

**Table 1.** Significant GO terms associated with “dark-turquoise” module, specific to conspecific song playback and “yellow” module, shared responses for conspecific song and cowbird chatter playback. FDR represents the *p*-value adjusted for false discovery rate. Significant GO terms associated with “yellow” module

**Significant GO terms associated with “yellow” module**
| term_ID | description | FDR |
| --- | --- | --- |
| GO:0051607 | defense response to virus | 3.16E-05 |
| GO:0060759 | regulation of response to cytokine stimulus | 0.006 |
| GO:0032480 | negative regulation of type I interferon production | 0.01 |
| GO:2000042 | negative regulation of double-strand break repair via homologous recombination | 0.04 |
| GO:0009607 | response to biotic stimulus | 0.04 |

**GO terms associated with “dark-turquoise” module**
| term_ID | description | FDR |
| --- | --- | --- |
| GO:0019219 | regulation of nucleobase-containing compound metabolic process | 3.90E-05 |
| GO:0044265 | cellular macromolecule catabolic process | 3.94E-05 |
| GO:0016567 | protein ubiquitination | 5.74E-05 |
| GO:0070647 | protein modification by small protein conjugation or removal | 6.24E-05 |
| GO:0044267 | cellular protein metabolic process | 0.001 |
| GO:0043412 | macromolecule modification | 0.003 |
| GO:0017148 | negative regulation of translation | 0.006 |
| GO:0032482 | Rab protein signal transduction | 0.03 |

## Discussion

Our expectations that subjects would sing in response to conspecific playback and approach, spend and perch close to redwing and cowbird playback were confirmed. We are therefore confident that we produced meaningful behavioral differences in aggressive responses of our subject male red-winged blackbirds. By studying free-living male territorial red-winged blackbirds, we detected patterns of co-expressed genes matching patterns of variation in aggressive behavioral responses to different classes of playbacks. In one behavioral response metric (approach distance) and in a co-expressed gene module, responses to conspecific songs and brood parasitic heterospecific calls were similar and both were significantly different from responses to harmless heterospecific calls. In turn, in a second behavioral response (number of songs) and another co-expressed gene module, responses to conspecific songs were significantly different from responses to both brood parasitic heterospecific and harmless heterospecific calls. Integrating the discriminability generated by these two patterns of responses provides for unique encoding for each of the three different playback types in both the behavioral and gene expression domains, separating conspecific songs from brood parasitic calls from harmless heterospecific calls.

Our results demonstrate that antagonistic responses to both conspecifics and brood parasites can involve similar physiological responses. The gene module correlated with both conspecific songs and cowbird chatters, relative to dove coos, was enriched for gene ontology immune system terms (Table 1), including defense responses to viruses and regulation of type I interferon production. For example, the top gene from the module, NLRC5 (NOD-like receptor family CARD domain containing 5 gene; Fig. 4), regulates adaptive immune responses against pathogens, including the activation of MHC class I genes ^21^. Unsurprisingly, these genes from the module were downregulated relative to the control playbacks, perhaps as a trade-off between aggressive responses and immune function seen elsewhere in avian and other vertebrate lineages^22^. Similarly, luteinizing hormone decreases in male red-wing blackbirds in response to simulated territorial intrusion ^23^. However, the source of the mRNA expression in peripheral blood, such as erythrocytes, leukocytes or exosomes ^24^, as well as the physiological function of these genes in either immunosuppressive or enhancement of immune response to acute stress remains unclear.

Overall, the behavioral and gene expression data indicate that similar physiological pathways may be involved in the cognitive recognition and motor responses to distinct antagonistic threats, in this case conspecific and heterospecific intruders. Many host species of brood parasites have evolved aggressive anti-parasitic behaviors ^6^, yet the cognitive and physiological mechanisms involved in this behavioral evolution remain unclear ^9^. Our study suggests that some of proximate responses to conspecific intruders were co-opted in the evolution of anti-parasitic aggression towards female cowbirds.

The gene expression responses found here in adult, territorial redwing males are different from the IEG patterns detected from juvenile redwing males, which showed that the only conspecific calls generated differential responses in the auditory forebrain relative to cowbird and dove calls ^10^. In turn, our behavioral data from adult red-winged blackbirds confirm previous findings in this and other avian hosts of brood parasitic species, which demonstrated behavioral responses to heterospecific parasitic models and auditory stimuli ^15,25–27^.

We did not find differential gene expression in a candidate set of potential biomarker genes, previously identified from peripheral whole blood in a conspecific vs. harmless heterospecific (dove) acoustic playback paradigm in captive female zebra finches ^20^. Given that our experimental treatments in the present study also included conspecific (redwing songs) vs. irrelevant heterospecific (dove) comparisons, we predicted that we would detect some of these same potential biomarker genes to be differentially expressed in free-living redwings. However, the use of different study species (zebra finches vs. red-winged blackbirds) or the different sexes of our subjects (female finches vs. male blackbirds) likely contributed to our inability to identify consistent gene expression differences within whole blood for conspecific recognition in songbirds.

In conclusion, our study demonstrates a parallel in behavioral and gene expression responses to simulated antagonistic threats. In particular, we find support for physiological responses involved in conspecific territorial aggression co-opted in the evolved recognition of heterospecific brood parasites. In addition, our study further demonstrates the utility for peripheral gene expression to study avian recognition and behavioral responses to social stimuli.

## Methods

### Study Species and Study Area

We studied red-winged blackbirds at Newark Road Prairie in Rock County, Wisconsin, USA (42° 32’ N, 89° 08’ W) during the breeding season of 2018. Newark Road Prairie is a 13-ha wet-mesic remnant prairie and sedge meadow habitat that supports about 35 male redwing territories ^28^. All males were captured in Potter traps baited with non-viable sunflower seeds and were banded with United States Geological Survey numbered aluminum bands and unique color combinations of plastic wraparound bands for individual identification (USGS permit # 20438 to KY); most females at this study site were not banded.

We used observations of territorial behaviors to place additional seed-baited traps within breeding territories of potential subjects and allowed them to use the traps without being captured for 1–3 weeks prior to playbacks. Previous work at this site on redwings’ responses to model cowbirds vs. harmless heterospecifics, coupled with their respective call playbacks, showed strongly graded aggressive responses to the former relative to the latter ^15^.

### Playbacks

We presented one of three playback types at each territory: (1) male conspecific songs (“Redwing”; highly salient), (2) female cowbird chatter calls (“Cowbird”; salient heterospecific vocalization of a brood parasite of redwing nests), or (3) dove coo (“Dove”; non-salient vocalizations of a harmless sympatric heterospecific) for broadcasts. Given that male red-winged blackbirds exhibit greater behavioral responses to female vs. male cowbird models ^15^, the chatter call, a specific call of female cowbirds was chosen to acoustically simulate brood parasite intrusion. To address pseudo-replication ^29^, one out of five available exemplars were assigned at random for each playback type. Audacity v 2.2.0 was used to filter playback stimuli above 2000 Hz and below 500 Hz, and normalize mean amplitude of all stimuli. Acoustic stimuli were matched in peak amplitude and duration.

Playback types and exemplar files were randomly assigned to territorial male redwings using a stratified balanced design to keep sample sizes per type similar. For each playback type we randomly chose one of the exemplars to broadcast from an iPhone 5 or 6 (Apple Inc., Cupertino, California, USA) connected to an Ecoxgear ECOXBT speaker (Grace Digital Audio, Peterborough, Ontario, Canada) via a 30-m auxiliary cable. Playbacks were broadcasted at 80–85 dB SPL at 1 m from the source (as measured by a sound pressure meter: Pyle PSPL01, Pyle Audio Inc., Brooklyn, New York, USA), which approximated natural amplitude (KY personal observations).

We used a 30 min. paradigm to induce (differential) gene expression (e.g., ^20^), but to avoid habituation in the field, each playback consisted of three 10-min segments for a total period of 30 min. The first and third segments were active sound broadcast periods and the second segment was a 10-min silent period. Each active period consisted of 10 1-min sub-segments in which an exemplar played at 0, 10, 20, 30, and 40 s, followed by 20 s of silence. As soon as the 30-min playback was completed, we removed the playback equipment and baited and set the trap; we aimed to capture each subject within 30 min of the end of the playback, banded all previously unmarked subjects as described above, took a blood sample of approximately 100 μL (see below), and released the subject. We recorded the time to capture for all samples obtained within the 30-min maximum.

Prior to playback, we placed two markers each 10 m from the speaker to facilitate measuring proximity time to the speaker within 10 m. During each 10-min segment we recorded (1) number of songs (songs), (2) number of flights towards the speaker (approaches), (3) time spent within 10 m of the speaker (time in proximity), and (4) the distance (m) of the closest approach (closest approach). These behavioral variables are well known to indicate a male redwing’s aggressiveness ^30^. We analyzed behavioral data collected during the 1st and 3rd 10-min periods, which preceded the capture timepoint by up to 30 min and, thus, is representative of the time sampled for the gene-expression patterns, too. Number of songs and number of approaches were the totals for periods 1 and 3 (active playback). Time (min) within 10 m was the total time for the active playback periods. Closest approach was the shortest distance between the subject and the speaker during the two active playback periods.

We used generalized linear mixed models with a negative binomial response and individual male identity as a random effect with glmmTMB in R (version 3.5.1) to analyze the responses of male redwings to the three playbacks. Each model included playback treatment, trail period, and whether the male was captured as explanatory variables. A significant result was further examined with a Tukey post-hoc analysis to identify significantly different pairs of treatment. The alpha level was set at the *p* < 0.05.

### RNA extraction and sequencing

A blood sample from each subject was collected from the brachial vein using a 1.27 cm 27 g needle and heparinized capillary tube (Fisher Scientific, Pittsburgh, Pennsylvania, USA). Blood was placed in 500 μl of RNAlater and then stored in a −80°C freezer for processing. RNA was extracted with RiboPure blood kits (Life Technologies, Carlsbad, California, USA) and treated with DNAse for purification. We assessed the quality of purified RNA on a Bioanalyzer (Agilent, Wilmington, Delaware, USA) (RIN > 7.0).

All library preparations and sequencing were performed at the University of Illinois at Urbana-Champaign Roy J. Carver Biotechnology Center, Urbana, IL, USA. A library for each sample was prepared with an Illumina TruSeq Stranded RNA sample prep kit. All libraries were pooled, quantitated by qPCR, and sequenced on one lane of an Illumina HiSeq 4000 with a HiSeq 4000 sequencing kit version 1, producing single-end 100 bp reads. Fastq files were demultiplexed with bcl2fastq v 2.17.1.14 (Illumina).

### Preparation of reference genome

Lacking an annotated reference genome for the red-winged blackbird, we created a proxy reference following the pseudo-it pipeline (https://github.com/bricesarver/pseudo-it). Briefly, we extracted DNA from liver and muscle tissue of a male red-winged blackbird (cataloged at the Museum of Southwestern Biology MSB:Bird:40979) and performed paired-end (200 bp) whole-genome sequencing on one lane of HiSeq (2500) at the Duke Genome Center. After removing the Illumina adapters with Trim Galore! v 0.3.7 (http://www.bioinformatics.babraham.ac.uk/projects/trim_galore/), which incorporates Cutadapt v 1.7.1 ^31^, we aligned the DNA reads to the closest related species publicly available, the white-throated sparrow (*Zonotrichia alibicolis*) genome (assembly Zonotrichia_albicollis-1.0.1), with BWA-mem ^32^. Using GATK ^33^ we then identified and inserted red-winged blackbird SNPs into the white-throated sparrow reference genome. To improve the proxy reference genome, we performed an additional iteration of the pseudo-it pipeline.

### Gene expression

We removed Illumina adapters from RNA reads with Trim Galore! v 0.3.7. We then aligned the reads to the proxy reference genome with Hisat2 ^34^ and quantified read abundance with HTSeq-count ^35^. With DeSeq2 ^36^, we then included the playback treatments and time to capture (minutes) to analyze the gene expression patterns of the 20 top genes identified from the parallel expression patterns of peripheral (whole blood) RNA-sequencing study of Louder et al. ^20^.

Next, we sought to identify networks of genes specifically responsive to Redwing song, Cowbird chatter or both Redwing and Cowbird relative to Dove coo. We performed weighted gene co-expression network analysis (WGCNA), which is used for finding clusters (modules) of highly correlated genes and determine the relationship of modules to treatments. We used the WGCNA package in R (Langfelder & Horvath 2008) to identify modules of co-expressed genes in our dataset. To remove genes with low read abundance, we filtered for genes with < 1 count per million in at least 10 samples. We then normalized for read-depth and extracted variance stabilizing transformed (vst) read counts from DEseq2 into WGCNA. To build the co-expression matrix, we chose a soft thresholding power (β) value of 12, at which at which we observed a plateau in Mean Connectivity, thus representing a scale-free topology ^37^. We generated a signed network with minimum module size of 30 genes and merged highly correlated modules (dissimilarity threshold = 0.25). We then correlated the eigengene, which is the first principal component of a module, of these merged modules with playback treatments (Redwing, Cowbird, Dove). Modules with *p* ≤ 0.05 were considered significantly correlated with a given trait.

Finally, we tested for functional enrichment of gene ontology (GO) categories with Gorilla ^38^. For each module, genes were ranked based on their module membership score determined in the WGCNA analysis. We preferred this rank-order based approach (as opposed to strict module assignment) as it reflects the correlation among modules, and because some genes could be assigned to multiple modules ^39^. GOrilla performs ranked-order analyses with human gene IDs, so we identified orthologous genes from the annotated red-winged blackbird proxy-genome. Statistical significance of GO categories was determined with *p*-values corrected for multiple hypothesis testing (*FDR* < 0.05). We used REVIGO to remove redundant and overlapping GO categories, with an allowed semantic similarity measure of 0.5 ^40^.

## Data Accessability

The data that support the findings of this study are being submitted to NCBI, and will be available prior to publication.

## Author contributions

M.I.M.L., K.Y., F.M.K.U. and M.E.H. designed the project; M.L., K.Y. collected data and samples; A.A.L. collected laboratory data, M.I.M.L., C.B. and M.E.H. analyzed the data; M.I.L, and M.E.H. wrote the first draft, and all authors provided critical feedback, reviewed, and edited the manuscript. All authors approve the manuscript.

## Acknowledgements

For financial support we thank the National Science Foundation IOS Program in Behavioral Systems (nos. 1456524 and 1456612 to M.E.H. and C.B.) and the National Academies Keck Futures Initiative Grant (to F.M.K.U. and M.E.H). Additional funding was provided by the Harley Jones Van Cleave Professorship at the University of Illinois (to M.E.H.) and the Mead-Witter Foundation (to K.Y.). For discussions, we are grateful to Kathleen Lynch, Mikus Abolins-Abols, and Sarah London. Capture and banding were conducted under USGS banding permit #20438 to K.Y. The playback protocol was approved by the Beloit College IACUC (protocol #18002).

